# TDP-43 regulates chromatin looping and gene transcription through binding and stabilizing DNA G-quadruplex structures

**DOI:** 10.64898/2026.05.13.724493

**Authors:** Feng Yang, Suyang Zhang, Xiaofan Guo, Yulong Qiao, Yuwei Zhang, Hao Sun, Xiaona Chen, Huating Wang

**Affiliations:** Department of Chemical Pathology, The Chinese University of Hong Kong, Hong Kong, China; Center for Neuromusculoskeletal Restorative Medicine Limited, Hong Kong Science Park, Hong Kong SAR, China; Department of Orthopaedics and Traumatology, The Chinese University of Hong Kong, Hong Kong, China; Li Ka Shing Institute of Health Sciences, The Chinese University of Hong Kong, Hong Kong SAR, China; Department of Bioinformatics, Warshel Institute for Computational Biology, Faculty of Medicine, Chinese University of Hong Kong (Shenzhen), Guangdong, China

**Keywords:** TDP-43, DNA G-quadruplexes, chromatin loops, transcriptional regulation

## Abstract

TAR DNA-binding protein 43 (TDP-43) is a multifunctional DNA/RNA-binding protein implicated in transcriptional and post-transcriptional regulation. Dysregulation of TDP-43 is closely correlated with human diseases such as cancer and neurodegenerative diseases. Although its roles in RNA metabolism are well characterized, its function in transcriptional regulation remains largely underexplored. DNA G-quadruplexes (dG4s) are non-canonical nucleic acid structures enriched at gene promoters and regulatory elements, where they facilitate chromatin looping and gene transcription. Here, we investigated the transcriptional regulatory role of TDP-43 by integrating multi-omics datasets, including Hi-C, dG4 ChIP-seq, TDP-43 ChIP-seq, RNA-seq and ATAC-seq from K562 and HepG2 cells. Our analyses demonstrate TDP-43 binding and dG4s formation are highly colocalized at chromatin loop anchors, particularly at promoter and enhancer regions. TDP-43 occupancy at these anchors correlates with increased dG4 stability, chromatin loop interaction frequency, elevated chromatin accessibility, and upregulated gene expression. Morover, TDP-43 knockdown in HepG2 cells revealed a significant reduction in dG4 formation and loop interaction strength, accompanied by widespread transcriptional dysregulation. Collectively, our findings highlight a novel regulatory role of TDP-43 in facilitating long-range chromatin interactions and transcriptional activation through binding to and stabilizing dG4 structures, providing a mechanistic basis for gene dysregulation driven by TDP-43 dysfunction in diseases.

## Introduction

TAR DNA-binding protein 43 (TDP-43) is a ubiquitously expressed nuclear protein that plays a pivotal role in gene regulation and RNA metabolism. Although initially identified as a DNA-binding protein that regulates the transcription of the HIV-1 gene, subsequent research has largely focused on its function as an RNA-binding protein (RBP) involved in regulating post-transcriptional RNA processing, including pre-mRNA splicing, mRNA transport, translation, and mRNA stability [1–3], with the broader role of TDP-43 in transcriptional regulation remaining underexplored. Dysregulation of TDP-43 profoundly contributes to various human diseases, including cancer and neurodegenerative diseases such as amyotrophic lateral sclerosis (ALS) and frontotemporal lobar degeneration (FTLD) [2,4]. It is thus imperative to elucidate the multifaceted regulatory roles of TDP-43 to establish a knowledge foundation for novel therapeutics.

TDP-43 regulates transcription through diverse mechanisms. It was first identified as a transcriptional repressor that binds the HIV-1 TAR element and modulates transcription complex assembly. TDP-43 can also directly bind to GTGTGT motifs to regulate RNA polymerase II pausing, affecting *acrv1* gene expression during spermatogenesis [5,6]. Subsequent genome-wide studies have revealed that TDP-43 localizes predominantly to promoters of actively transcribed genes, where it influences nascent RNA synthesis without direct association with RNA polymerase II and plays a critical role in maintaining the transcriptional stability of protein-coding genes and transposable Alu elements [7]. Moreover, recent studies have revealed that TDP-43 can also regulate gene transcription through binding DNA at specific motifs and regulatory elements to modulate higher-order genome structures, particularly enhancer-promoter (E-P) interactions, which are pivotal for precise transcriptional control. The Drosophila TDP-43 homolog TBPH has been shown to recruit cohesin to distal regulatory elements via GU-rich motifs in nascent transcripts [8]. In mammalian cells, TDP43 has been shown to regulate long-range regulatory interactions and epigenetic crosstalk to modulate gene transcription. Loss of TDP-43 depletes R-loops and 5-hydroxymethylcytosine (5hmC) within gene bodies and enhancers of downregulated genes, disrupting E-P looping and repressing gene transcription [9]. However, the mechanisms underlying TDP-43-dependent chromatin architecture regulation are not fully understood.

Interestingly, a recent study has revealed that TDP-43 can interact with and stabilize DNA G-quadruplexes (G4s) *in vitro* [10].. G4s are non-canonical secondary structures formed in single-stranded guanine-rich DNA or RNA sequences, typically consisting of four tracts of guanines align in stacked tetra planes stabilized by Hoogsteen hydrogen bonding [13,14]. DNA G4s (dG4s) participate in regulating a wide range of biological processes, including transcription, telomere maintenance, DNA replication etc. Notably, recent advances in genome-wide mapping of endogenous dG4 formation have revealed the enrichment of dG4s at gene regulatory regions such as promoters and enhancers and the important roles of dG4 in regulating gene transcription. Moreover, dG4 structures are increasingly recognized as dynamic platforms that recruit specific proteins to facilitate long-range chromatin interactions and modulate transcriptional regulation [11,12,15]. For example, our recent study has shown that dG4s interact with transcription factor (TF) MAX to regulate gene transcription in skeletal muscle stem cells [15]. Nevertheless, the interplay between TDP-43 and dG4s in transcriptional regulation remains to be explored.

In this study, we integrated multi-omics datasets, including dG4 ChIP-seq, TDP-43 ChIP-seq, Hi-C, RNA-seq, ATAC-seq, etc., from K562 and HepG2 cells to investigate the interplay between TDP-43 and dG4 in regulating gene transcription. We demonstrate that TDP-43 and dG4s are highly colocalized and enriched at chromatin loop anchors genome-wide. TDP-43 binding at chromatin loop anchors is associated with higher dG4 enrichment and increased interaction frequency, accompanied by increased chromatin openness, TF binding, and gene expression. TDP-43 knockdown significantly diminishes dG4 signals, reduces loop interaction frequency, and leads to widespread transcriptional dysregulation. Altogether our findings highlight a novel regulatory role for TDP-43 in stabilizing dG4 at chromatin loop anchors to facilitate looping and associated gene transcription.

## Results

### Integrated multi-omics analyses reveal extensive co-occupancy of TDP-43 and dG4s at gene promoters in K562 and HepG2 cells

To investigate the potential functional interplay between TDP-43 and dG4 in transcriptional regulation, we integrated publicly available multi-omics datasets from K562 (chronic myelogenous leukemia) and HepG2 (hepatocellular carcinoma) cells [16–18], including TDP-43 ChIP-seq, dG4 ChIP-seq, Hi-C-seq, ATAC-seq, RNA-seq, and TFs ChIP-seq datasets (Fig. 1A) . Re-analysis of these datasets identified 8,789 and 7,511 TDP-43 binding sites in K562 and HepG2 cells, 9,795 and 8,803 dG4 sites were also identified (Supplementary Table S1). Interestingly, both TDP-43 and dG4 sites were predominantly enriched at promoter and distal intergenic regions in both cells (Fig. 1B-C). In K562 and HepG2 cells, promoter regions accounted for 75.64%/81.96% of TDP-43 and 82.79%/75.52% of dG4 binding sites, while distal intergenic regions contained 10.26%/7% and 7.95%/8.45%. Notably, 2,908 (33.09%) of the total TDP43 peaks in K562 cells and 2,087 (27.79%) in HepG2 cells overlapped with the dG4 peaks, with the vast majority of these overlapping peaks located in promoter regions (Fig. 1D-E). A closer examination at the protein-coding gene promoter regions revealed that 2,836 of the 4,425 TDP-43-bound promoters (64.09%) in K562 cells and 2,508 of the 4,570 TDP-43-bound promoters (54.88%) in HepG2 cells harbored dG4 structures (Fig. 1F, Supplementary Table S2). Representative genomic loci illustrating the co-occupancy of TDP-43 binding and dG4 include the *TAL1* (TAL BHLH Transcription Factor 1 gene) promoter in K562 cells which is essential for the development of all hematopoietic lineages [19,20], and the *DBF4* (DBF4-CDC7 Kinase Regulatory Subunit) promoter in HepG2 cells, which is critical for DNA replication [21,22] (Fig. 1G-H). Collectively, these findings demonstrate extensive genome-wide co-localization of TDP-43 binding and dG4 structures, particularly at gene promoters, suggesting a potential synergistic role of TDP-43 and dG4 in regulating gene transcription.

**Figure 1.**
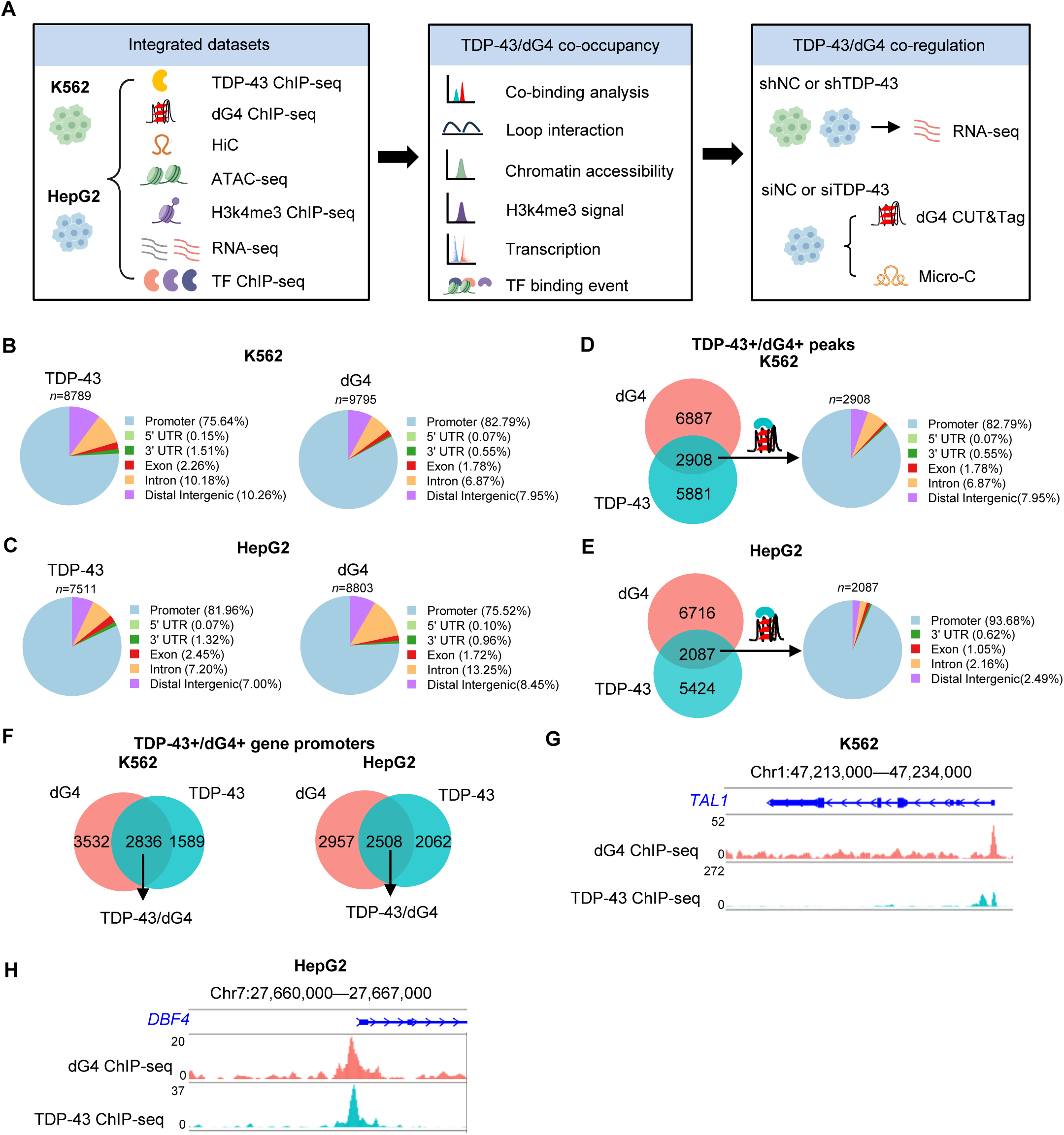
Integrated multi-omics analyses reveal extensive co-occupancy of TDP-43 and dG4s at gene promoters in K562 and HepG2 cells. **(A)** Schematic of the overall design of the study. **(B-C)** Genomic location profiling of TDP-43 and dG4 in K562 and HepG2 cell lines. **(D-E)** Genomic location profiling of sites with TDP-43 and dG4 co-occupancy. **(F)** Overlap of genes with promoter TDP-43 binding and genes with promoter dG4 formation in K562 and HepG2 cells. **(G-H)** Genome browser view of dG4 and TDP-43 signals across the promoters of *TAL1* in K562 cells, and *DBF4* in HepG2 cells.

### TDP-43-mediated transcriptional regulation correlates with dG4 co-occupancy

To examine whether the regulatory role of TDP-43 on transcription was associated with dG4, we analyzed publicly available RNA-seq data from K562 and HepG2 cells with TDP-43 knockdown by shRNA (Fig. 2A). The knockdown resulted in 15,972 and 10, 072 differentially expressed genes (DEGs) in K562 and HepG2 cells (Supplementary Fig. S1A-F), among which 3,262 and 2,305 were directly bound by TDP-43 at their promoters (Supplementary Fig. S1G-L). Notably, 2,126 (65.17%) of the TDP-43-bound DEGs in K562 cells and 1,305 (56.62%) in HepG2 cells also harbored promoter dG4s (Fig. 2B, Supplementary Table S3), constituting as possible TDP-43/G4 co-regulated transcriptional targets. Among them, 894 were up-regulated, and 1,232 were down-regulated in K562 cells, while 616 were up-regulated and 689 were down-regulated in HepG2 cells (Fig. 2C, Supplementary Table S3). Pathway enrichment analysis revealed distinct functional signatures for these genes. In K562 cells, up-regulated TDP-43/dG4 targets were enriched in chromosome segregation and double-strand break repair (such as *KIF4*), whereas down-regulated targets were associated with RNA splicing and cytoplasmic translation (such as *EIF2D*) (Fig. 2D and F, Supplementary Table S3). In HepG2 cells, up-regulated targets were enriched for intracellular transport and protein catabolic process (such as *CDC42*), while the down-regulated targets were associated with DNA replication (such as *MCM4*) (Fig. 2E and G, Supplementary Table S3). Altogether, these data demonstrate that the transcriptional regulatory function of TDP-43 may be largely mediated in a dG4-dependent manner.

**Figure 2.**
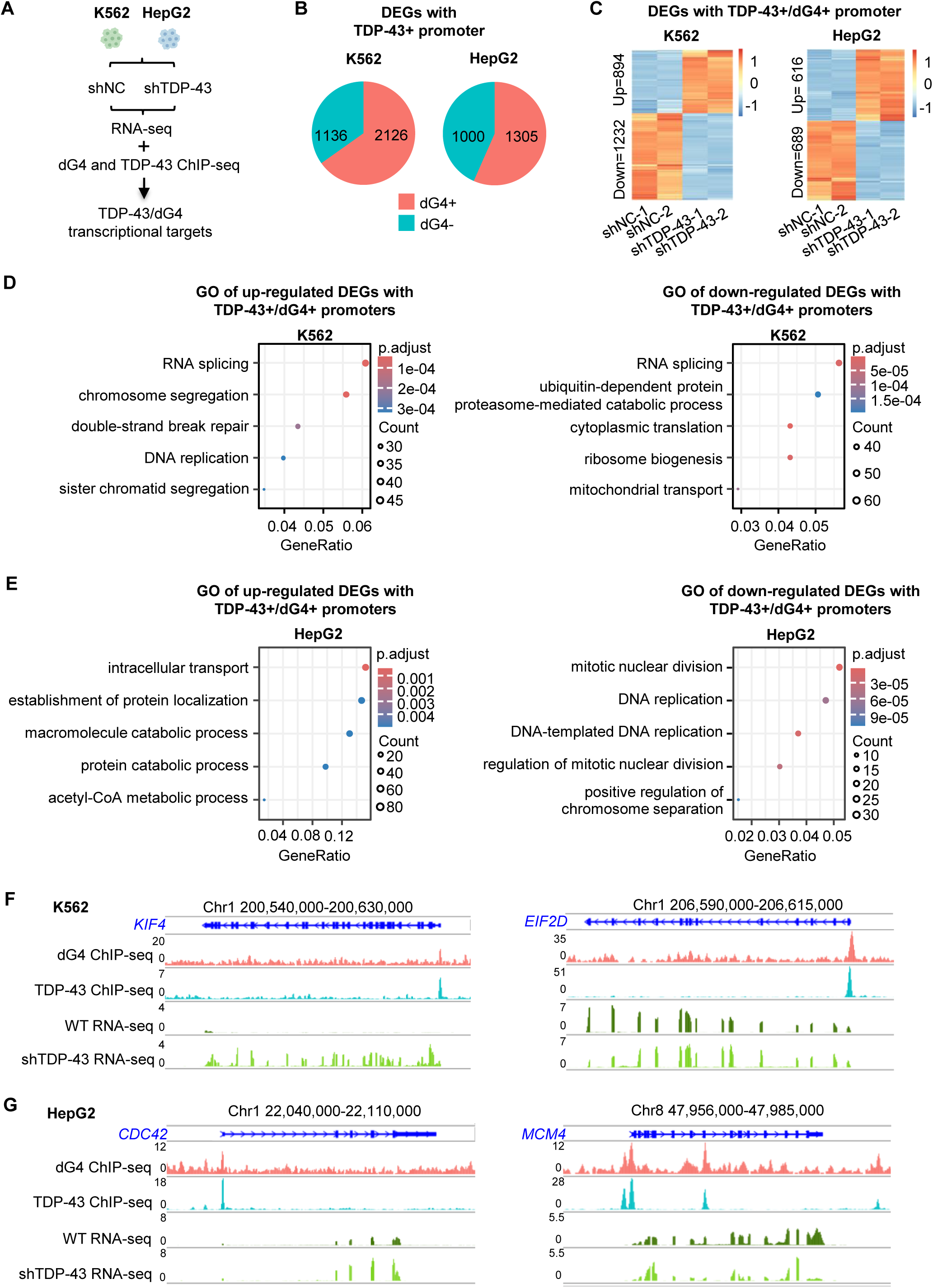
TDP-43-mediated transcriptional regulation correlates with dG4 co-occupancy. **(A)** Schematic of the analysis combining RNA-seq datasets in K562 and HepG2 cells with shNC or shTDP-43 knockdown and dG4 and TDP-43 ChIP-seq datasets. **(B)** Pie charts of DEGs with promoter TDP-43 binding in shNC vs shTDP-43 K562 and HepG2 cells overlapping with promoter dG4 peaks. **(C)** Heatmaps of DEGs with promoter TDP-43/dG4 in K562 and HepG2 cells. **(D-E)** GO term analysis of up-regulated and down-regulated DEGs with promoter TDP-43/dG4 targets in K562 and HepG2 cells. **(F-G)** Snapshots of TDP-43 and dG4 signals across *KIF4, EIF2D* promoters in K562 cells and *CDC42*, *MCM4* promoters in HepG2 cells.

### TDP-43 binding is associated with dG4 stabilization at chromatin loop anchors

To further elucidate the TDP-43/dG4 co-regulatory mechanism, we examined whether TDP-43/dG4 were involved in regulating chromatin loops. Using publicly available Hi-C data, we identified a total of 13,621 and 14,513 high-confidence chromatin loops in K562 and HepG2 cells, respectively. A robust accumulation of TDP-43 and dG4 signals at loop anchors was observed compared to the flanking genomic regions (Fig. 3A-B, Supplementary Table S4). Notably, approximately 50% of TDP-43-bound loop anchors (757 in K562; 962 in HepG2) were co-occupied by dG4s (Fig. 3C). Moreover, TDP-43 and dG4 signal intensities were positively correlated at these co-occupied anchors (Fig. 3D). Moreover, we found that the TDP-43+/dG4+ loop anchors exhibited a significantly higher dG4 signal than TDP-43–/dG4+ loop anchors (Fig. 3E-F), which were exemplified on the *SF3B5* locus in K562 cells and *COMMD3* locus in HepG2 cells (Fig. 3G). Altogether, the above results indicate that TDP-43 may bind and stabilize dG4 at chromatin loop anchors.

**Figure 3.**
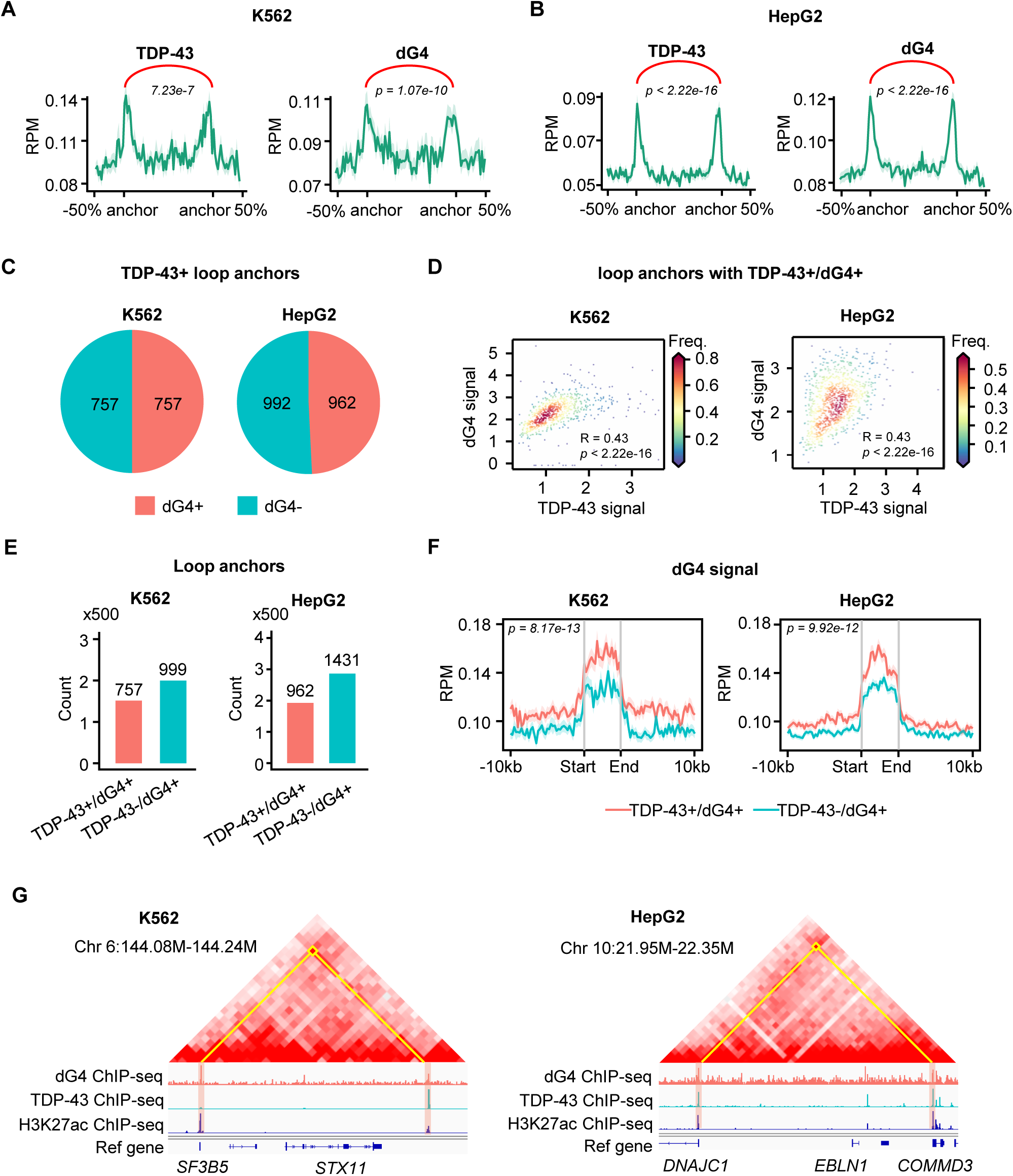
TDP-43 binding is associated with dG4 stabilization at chromatin loop anchors. **(A)** TDP-43 and dG4 signal comparison of loop anchors and surrounding regions in K562 cells, the statistical significance was calculated by Mann-Whitney U test, p = 7.23 × 10^-07^ for TDP-43 signal, p =1.07 × 10^-10^ for dG4 signal. **(B)** TDP-43 and dG4 signal comparison in HepG2 cells, the statistical significance was calculated by Mann-Whitney U test, p < 2.22 × 10^-16^ for TDP-43 signal and dG4 signal. **(C)** TDP-43 bound loop anchors overlaps with dG4 in K562 and HepG2 cells. **(D)** Correlation of TDP-43 and dG4 signal at TDP-43+/dG4+ loop anchors in K562 and HepG2 cells. **(E)** Number of loop anchors with dG4 formation overlapping with TDP-43 in K562 and HepG2 cell lines. **(F)** Comparison of dG4 signal at dG4-formed loop anchors with or without TDP-43 binding, the statistical significance was calculated by Mann-Whitney U test. **(G)** Genome browser view of TDP-43, dG4 and Hi-C signals across the promoters of *SF3B5* (K562 cell) and *COMMD3* (HepG2 cell).

### TDP-43/dG4 co-occupancy facilitates chromatin looping and promotes gene transcription

To examine whether TDP-43 /dG4 co-occupancy on loop anchors may play a role in chromatin looping and gene transcription, we first classified the loops into four types according to TDP-43 binding and dG4 formation at the loop anchors: TDP-43+/dG4+ (at least one loop anchor with both TDP-43 and dG4), TDP-43+/dG4- (at least one loop anchor with TDP-43 and none with dG4), TDP-43-/dG4+ (at least one loop anchor with dG4 and none with TDP-43) and TDP-43-/dG4- (neither anchors with TDP-43 or dG4) loops (the loops with dG4 at one anchor contain G4, and TDP-43 at the other were excluded from subsequent analysis to maintain the mutual exclusivity of the four defined categories). (Fig. 4A, Supplementary Fig. S2A, and Supplementary Table S5). In both K562 and HepG2 cells, TDP-43+/dG4+ loops exhibited the highest chromatin interaction frequency compared with other types, while TDP-43+/dG4- and TDP-43-/dG4+ loops showed similar interaction frequency, which were both significantly higher than the TDP-43-/dG4-loops (Fig. 4B). Moreover, genes associated with TDP-43+/dG4+ loop anchors (Fig. 4C) also showed significantly higher expression levels compared to other types (Fig. 4D). In line with the findings, promoters associated with TDP-43+/dG4+ loop anchors exhibited significantly higher ATAC-seq and H3K4me3 ChIP-seq signals, as well as TF binding events compared to other loop anchors (Fig. 4E-G, Supplementary Table S6). Altogether, the above findings demonstrate that TDP-43 and dG4 co-binding on loop anchors may synergistically enhance chromatin looping and promoting gene expression.

**Figure 4.**
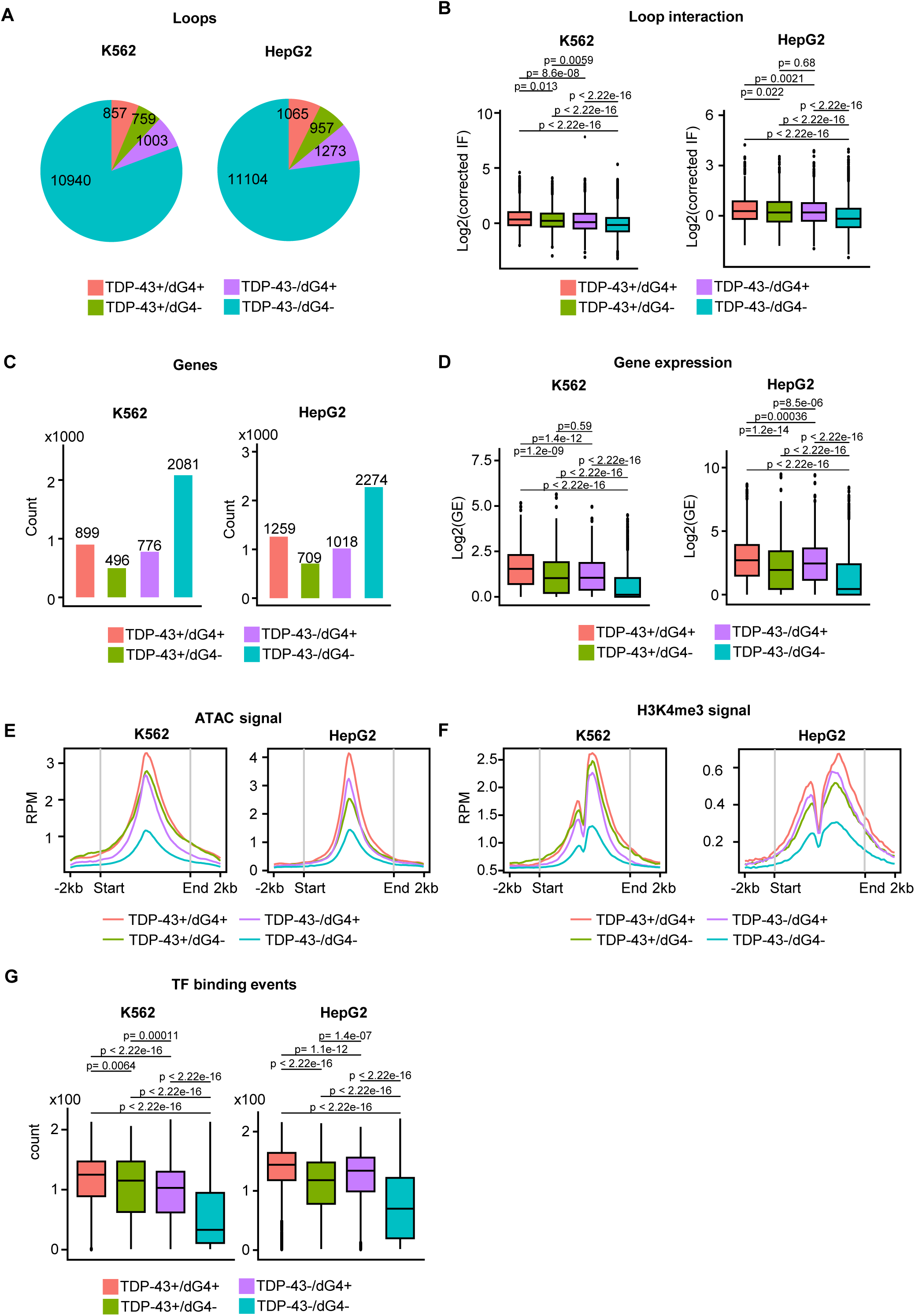
TDP-43/dG4 co-occupancy facilitates chromatin looping and promotes gene transcription. **(A)** Number of TDP-43+/dG4+, TDP-43+/dG4-, TDP-43-/dG4+ and TDP-43-/dG4-loops in K562 and HepG2 cells. **(B)** Interaction frequency comparison of above four types of loops in K562 and HepG2 cells. **(C)** Count of genes whose promoter located at the above four type loop anchors in K562 and HepG2 cells. **(D)** Expression level of the genes in **(C)** in K562 and HepG2 cells. **(E)** Comparison of ATAC-seq signal, **(F)** H3K4me3 signal, and **(G)** TF binding events of gene promoters in **(C)** in K562 and HepG2 cells. The statistical significance in (B), (D), (E), (F), (G) was calculated by Mann-Whitney U test, p-values were shown in the figures.

### TDP-43/dG4 co-occupancy regulates target genes with crucial biological functions

To further substantiate the functional role of TDP-43/dG4 co-occupancy, we analyzed the genes associated with TDP-43+/dG4+ loop anchors. In K562 cells, the genes were mainly related to RNA processing associated terms, e.g., RNA processing and mRNA processing; In HepG2 cells, these genes were enriched for cell cycle associated terms, e.g., cell cycle and cell cycle process (Fig. 5A-B), suggesting TDP-43/dG4 co-occupancy regulates genes with essential biological functions. Consistently, TDP-43 knockdown resulted in the highest proportion of down-regulated DEGs associated with the TDP-43+/dG4+ loop anchors compared with other types of loop anchors in both K562 and HepG2 cells (Fig. 5C-D). These down-regulated genes were again enriched for RNA processing associated terms in K562 cells, and cell cycle associated terms in HepG2 cells (Fig. 5E-F). The above results suggested that TDP-43/dG4 co-occupancy at loop anchors facilitates chromatin looping to regulate a group of target genes with crucial biological functions in both K562 and HepG2 cells.

**Figure 5.**
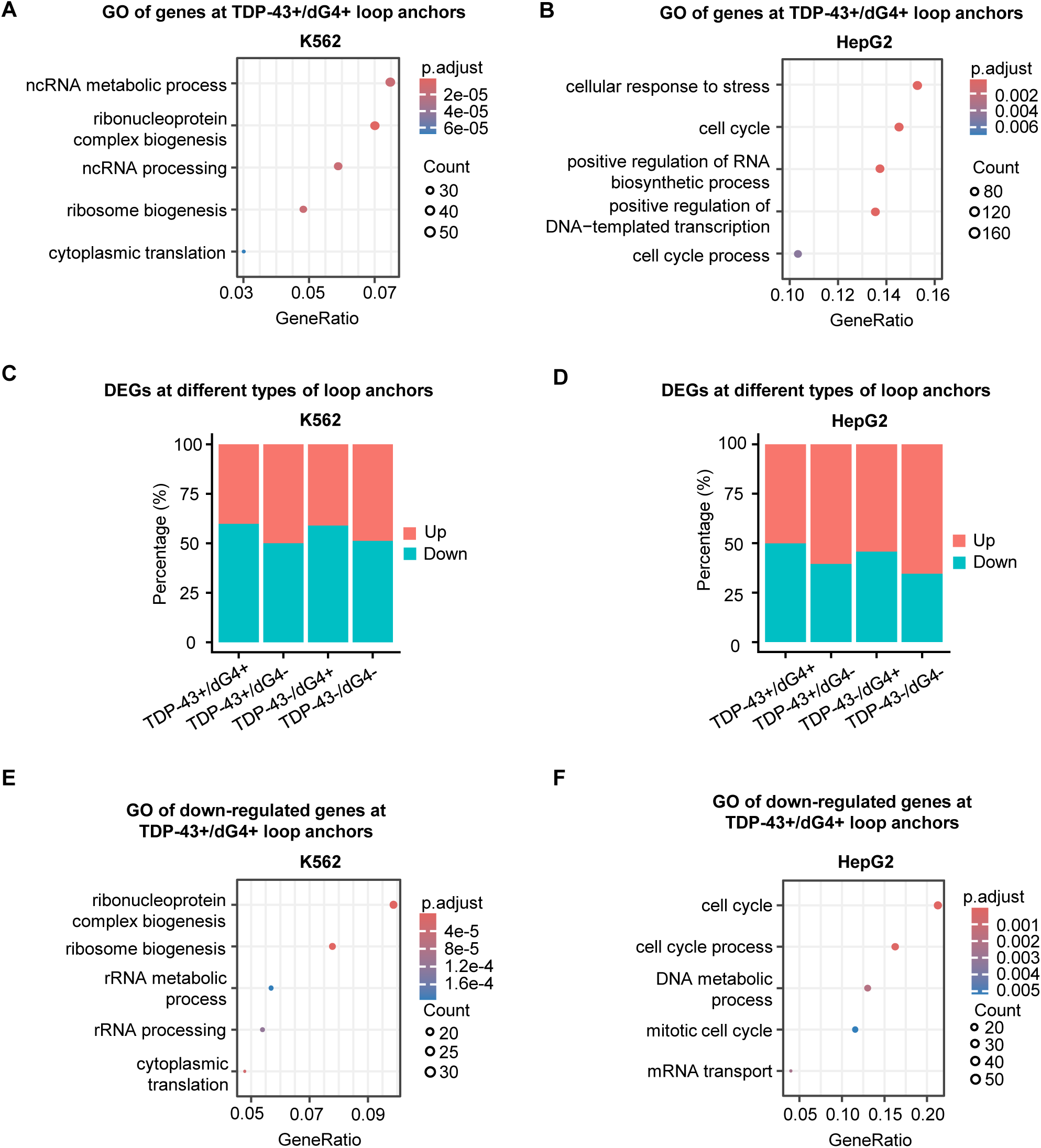
TDP-43/dG4 co-occupancy regulates target genes with crucial biological functions. **(A)** GO term enrichment analysis of genes at dG4+/TDP-43+ loop anchors in K562 and (**B**) HepG2 cells. **(C)** The proportion of up- or down-regulated genes at different types of loop anchors in K562 and (**D**) HepG2 cells. **(E)** GO term enrichment analysis of down-regulated genes at dG4+/TDP-4+ loop anchors in K562 and (**F**) HepG2 cells.

### TDP-43 stabilizes dG4 formation at loop anchors in HepG2 cells

To experimentally validate our hypothesis that TDP-43 stabilizes dG4 to modulate chromatin looping and gene transcription, we knocked down TDP-43 expression in HepG2 cells with siRNA oligos (Fig. 6A-6B). dG4 CUT&Tag was conducted to map the impact of TDP-43 loss on dG4 formation. The three biological replicates showed high reproducibility (Pearson correlation coefficient, r > 0.8) within groups (Supplementary Fig. S3A). In total, we identified 4,077 and 3,024 dG4 peaks in siNC and siTDP-43 groups, respectively (Fig. 6C, Supplementary Table S7). Motif scanning revealed a high prevalence of G-rich sequences in the dG4 peaks (Supplementary Fig. S3B), and the majority (89% in siNC and 90% in TDP-43) overlapped with *in vitro* identified potential dG4-forming sequence (PQS) (Supplementary Fig. S3C). Consistent with previous dG4 ChIP-seq data, dG4 CUT&Tag signals were primarily enriched at promoters (89.49% and 91.69%) and distal intergenic regions (4.15% and 3.34%) (Fig. 6D). A comparative analysis revealed that while 62.05% of the dG4 peaks remained unaltered, TDP-43 loss induced a clear reduction of both total dG4 peaks number and signal intensity (Fig. 6E). Notably, at the TDP-43 binding sites, dG4 signal intensity significantly decreased upon TDP-43 knockdown (Fig. 6F), supporting the hypothesis that TDP-43 binding stabilizes dG4 formation in HepG2 cells.

**Figure 6.**
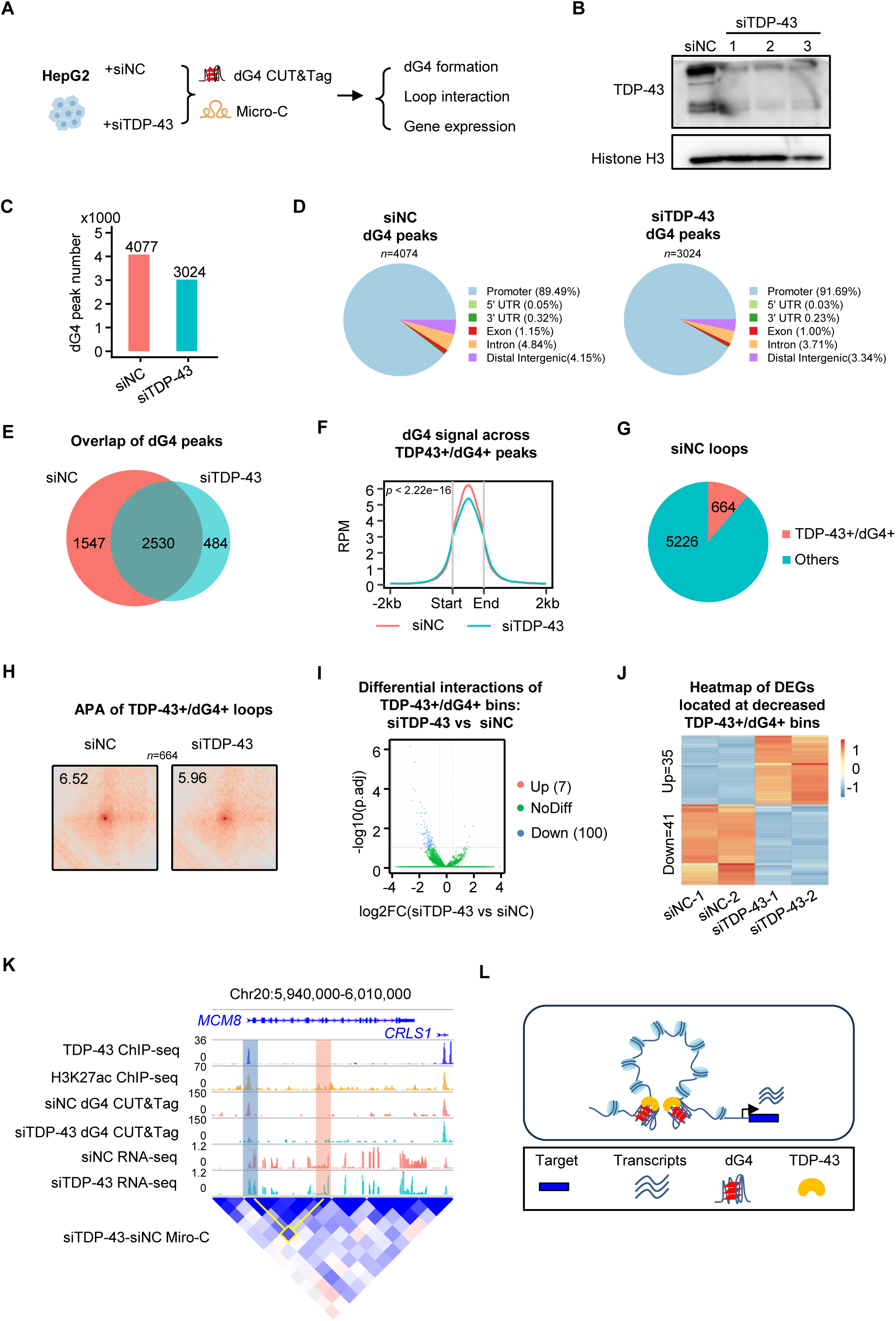
TDP-43 stabilizes dG4 formation at loop anchors in HepG2 cells. **(A)** Scheme of dG4 CUT&Tag and Micro-C sequencing of HepG2 cells transfected with siNC or siTDP-43 oligos. **(B)** Western blot of TDP-43 with HepG2 cells transfected with siNC or siTDP-43 oligos. H3 was used as the loading control. **(C)** dG4 peaks in HepG2 cells transfected with siNC or siTDP-43 oligos. **(D)** Genome location profiling of dG4 peaks in HepG2 cells transfected with siNC or siTDP-43 oligos. **(E)** Overlapping of dG4 peaks identified in HepG2 cells transfected with siNC or siTDP-43 oligos. **(F)** dG4 signals across TDP43/dG4 peaks in HepG2 cells transfected with siNC or siTDP-43 oligos; the statistical significance was calculated by Mann-Whitney U test, p < 2.2 × 10^-16^. **(G)** Number of TDP-43+/dG4+ loops and other loops identified by Micro-C in siNC HepG2 cells. **(H)** APA score of TDP-43+/dG4+ loops in (H) in HepG2 cells transfected with siNC or siTDP-43 oligos. **(I)** Volcano plot of differential interactions of TDP-43+/dG4+ bins in siTDP-43 vs siNC HepG2 cells. **(J)** Heatmap plot of DEGs located at decreased TDP-43+/dG4+ bins in siTDP-43 vs siNC HepG2 cells. **(K)** Snapshot of TDP-43 ChIP-seq, H3K27ac ChIP-seq, dG4 CUT&tag, RNA-seq and Micro-C signals across the *MCM8* promoter in HepG2 cells transfected with siNC or siTDP-43 oligos. **(L)** Schematic illustration of the model that TDP-43 loss disrupted dG4 formation, impairs chromatin looping and gene transcription.

We then conducted Micro-C which enables nucleosome resolution chromosome loop profiling. A total of 5,890 and 2,485 loops were identified in siNC vs siTDP-43 HepG2 cells by *Hiccups* (Supplementary Table S8). By integrating TDP-43 ChIP-seq data and the dG4 CUT&Tag-seq data, 664 TDP43+/dG4+ loops were identified in siNC and their interaction strength was significantly decreased with TDP-43 knockdown (APA 6.52 vs 5.96) (Fig. 6G-H). Differential interaction analysis using *HiCcompare* revealed that siTDP-43 knockdown resulted in 100 significantly weakened interactions compared to only 7 strengthened interactions (Fig. 6I). Consistently, RNA-seq analysis identified 41 down-regulated and 35 up-regulated genes associated with the disrupted looping (Fig. 6J). A representative example is *MCM8,* an important regulator for multiple DNA-related processes and pathologies [23], which exhibited robust TDP-43/dG4 co-occupancy at its promoter in siNC cells. Upon TDP-43 knockdown, a concomitant reduction of the dG4 signal, promoter-associated loop interaction frequency, and *MCM8* expression were observed (Fig. 6K). Altogether, the above findings indicate that TDP-43 binding stabilizes dG4 formation at chromatin loop anchors and synergistically facilitates chromatin loop interactions to promote gene expression.

## Discussion

In this study, we uncovered a previously uncharacterized function of TDP-43 in regulating chromatin looping and gene transcription through binding and stabilizing dG4s (Fig. 6L). Our integrative multi-omics analyses reveal that TDP-43 acts as a chromatin-associated factor that colocalizes with dG4s at loop anchors, particularly at promoter and enhancer regions, and this co-occupancy is associated with increased loop interaction frequency, chromatin accessibility, active transcription histone modification, TF binding, and gene expression. Consistently, TDP-43 knockdown in HepG2 cells led to a reduction in dG4 signals and loop interaction strength, resulting in transcriptional dysregulation. Our findings, in sum, demonstrate that TDP-43-mediated dG4 stabilization facilitates chromatin loop interactions, thereby maintaining a permissive chromatin environment for active transcription.

TDP-43 has been extensively characterized as an RBP that regulates RNA splicing, transport, and stability, etc. [1–3], while its potential role in regulating chromatin architecture remains largely unexplored. Our data reveals a novel function of TDP-43 in transcriptional regulation through its enrichment at promoter-enhancer loop anchors genome-wide and the occupancy is associated with elevated chromatin loop interaction frequencies, active histone modifications, TF binding, and gene expression in HepG2 and K562 cells. Notably, TDP-43 knockdown in HepG2 cells resulted in a genome-wide reduction in chromatin loop interaction strength and widespread transcriptional dysregulation, indicating that TDP-43 actively regulates loop architecture rather than occupying anchors passively. Our findings complement a recent study demonstrating that TDP-43 knockdown disrupts long-range E-P interactions and downregulates distal gene expression through depleting R-loops and 5-hydroxymethylcytosine (5hmC) at enhancers in SH-SY5Y neuroblastoma cells [9], revealing the role of TDP-43 as an important regulator of chromatin looping and transcription regulation with a novel DNA-centric mechanism.

Mechanistically, we showed that the transcriptional regulatory function of TDP-43 is tightly associated with dG4s. TDP-43 has been shown to bind to RNA G4 structures to regulate mRNA stability and translation [24], and RNA G4 in turn can modulate TDP-43 aggregation [25]. Recent *in vitro* studies have shown that TDP-43 can interact with dG4 formed at *cMyc* promoter [10]. Our work provides the first *in vivo* evidence that TDP-43 binding is associated with dG4 formation in both HepG2 and K562 cells. In HepG2 cells, we performed loss of function study to demonstrate that TDP-43 depletion reduced dG4 signals concomitantly with loss of loop interaction strength, supporting a model in which dG4s function as structural scaffolds and binding hubs for TFs, architectural proteins, and chromatin remodeling complexes. By stabilizing these dG4 scaffolds, TDP-43 couples local DNA secondary structure to modulate higher-order chromatin organization and transcriptional control, expanding the functional spectrum of both TDP-43 and dG4 biology. Nevertheless, since approximately half of TDP-43 binding sites at loop anchors lack detectable dG4s, other dG4-independent mechanisms are likely at play, e.g., protein-protein interactions with architectural factors or modulation of alternative DNA structures, merit exploration to fully delineate the role of TDP-43 in modulating 3D genome structure and gene transcription.

In cancer cells, alterations in 3D genome organization, including aberrant chromatin looping, enhancer hijacking, and disruption of topologically associating domains (TADs), are well-recognized drivers of oncogenic transcriptional programs [26]. While TDP-43 is frequently overexpressed in multiple solid tumors and has been linked to enhanced proliferation, apoptosis resistance, and poor prognosis [27–29], current knowledge of its oncogenic effects have been mainly correlated to its activity as an RNA-binding protein (RBP), including regulation of alternative splicing (e.g., *PAR3* and *NUMB* in breast cancer cell) and stability of specific mRNAs involved in metabolism and survival. Our findings reveal a novel DNA-centric mechanism that TDP-43 may contribute to gene dysregulation in cancer cells by modulating dG4-anchored chromatin loops. Notably, the regulatory targets of TDP-43 exhibited cancer-type specificity, as they are mainly enriched for cell cycle regulation in HepG2 RNA processing in K562 cells, suggesting that TDP-43 may regulate distinct target genes depending on cellular context. Moreover, given the important role of TDP-43 in neurodegenerative disease such as ALS and FTLD, our findings raise the intriguing possibility that nuclear depletion or mislocalization of TDP-43 in neurodegenerative disease may also compromise chromatin loop integrity and contribute to the transcriptional dysregulation observed in affected neurons, which can be investigated in future studies.

In conclusion, our study uncovers TDP-43 as a critical regulator of 3D genome organization and transcriptional control through direct binding and stabilization of dG4. These findings expand our understanding of TDP-43 as a multifunctional gene regulatory factor and open new therapeutic avenues for diseases driven by TDP-43 dysfunction.

## Methods

### Cell culture and transfection

HepG2 cells were obtained from the American Type Culture Collection (ATCC). Cells were maintained in medium consisting of DMEM supplemented with 10% FBS, 100 units/ml penicillin, and 100 μg/ml streptomycin, and cultured at 37°C in an atmosphere with 5% CO₂. HepG2 cells were transfected with siNC or siTDP-43 oligos (GenePharma, siRNA sequences provided in Supplementary Table S11) using Lipofectamine 3000 with a final concentration of 100 nM. 48h after transfection, the cell medium was discarded and the cells were collected for further analysis.

### Western blot

The cells were lysed by RIPA buffer with proteinase inhibitor (Thermo Fisher Scientific, 88266) and incubated on ice for 15 min. Then the cell lysis was centrifuged at 12000g for 10 min and the supernatant was used to conduct the western blot. The protein samples were then loaded to SDS-PAGE and the PVDF membrane with proteins was blocked by 3% BSA. The following dilutions were used for each antibody: TDP-43 (ABclonal, A1183, 1:1000), H3(Santa Cruz Biotechnology, sc-517576, 1:500); and the secondary antibodies: HRP-conjugated Goat anti-Rabbit IgG (ABclonal, AS014, 1:2000), HRP-conjugated Goat anti-Mouse IgG (ABclonal, AS003, 1:2000). Protein expression was visualized using an enhanced chemiluminescence detection system (GE Healthcare, Little Chalfont, UK).

### CUT&Tag

CUT&Tag assay was conducted using 100,000 HepG2 cells (Transfected by siNC or siTDP-43) with the CUT&Tag assay kit (Cell Signaling Technology, 77552). In brief, HepG2 cells were harvested and washed by cell wash buffer, then bound to concanavalin A-coated magnetic beads. Digitonin Wash Buffer was used for permeabilization. After that, cells were incubated with 2 μg of BG4 (ImmunoDiagnostics Limited), H3K4me3 (Cell Signaling Technology, 9751) or H3K27ac (Cell Signaling Technology, 8173T) antibody overnight at 4 °C. For dG4 CUT&Tag, an extra incubation step with 2 μg of anti-flag antibody (Sigma, F1804) was performed at room temperature for 1 hour, followed by the regular incubation step with 1ul of Secondary antibody (Cell Signaling Technology, Goat Anti-Rabbit IgG (H+L) Antibody #35401 or Donkey Anti-Mouse IgG (H+L) Antibody #52885) at room temperature for 30 minutes. Then, cell-bead slurry was washed with Digitonin Wash Buffer and incubated with Protein AG-Tn5 for 1 h at room temperature. After washing with High Salt Digitonin Buffer, Tagmentation Buffer was added into the cell-bead slurry to initiate Protein AG-Tn5 digestion, which was then incubated at 37°C for 1 hour. Then 0.5 M EDTA (Cell Signaling Technology, 7011), 10% SDS (Cell Signaling Technology, 20533) and 20 mg/mL Proteinase K (Cell Signaling Technology, 10012) were added to the reaction to stop the digestion. CUT&Tag fragments were released by incubation for 1 hour at 58 °C followed by centrifugation. After centrifugation, the supernatant was recovered, and DNA purification was performed by using DNA Purification Columns (Cell Signaling Technology). For DNA library construction, a CUT&Tag Dual Index Primers and PCR Master Mix for Illumina (Cell Signaling Technology, 47415) was used according to the manufacturer’s instructions. Bioanalyzer analysis and qPCR were used to measure the quality of DNA libraries including the DNA size and purity. Finally, DNA libraries were sequenced on the GeneMind SURFSeq 5000 platform.

### Micro-C

Micro-C was performed following the published protocol for mammalian Micro-C [30]. Cells were first crosslinked at 1 ml per million cells of 3 mM DSG crosslinker (MedChemExpress, HY-114697) for 35 min at room temperature, and then 1% formaldehyde was added and incubated for 10 more minutes. The crosslink was quenched with 0.375 M Tris (pH 7.0) for 5 min. Cells were pelleted at 1000 g at 4 °C for 5 min and resuspended with ice-cold PBS. One million cells were split into each tube and pelleted by centrifugation. After snap frozen in liquid nitrogen, cells were kept at − 80 °C. For cell lysis, cells were thawed on ice for 5 min, and lysed with 0.5 ml MB 1 buffer (10 mM Tris–HCl, pH 7.5, 50 mM NaCl, 5 mM, MgCl2, 3 mM CaCl2, 0.2% NP-40, 1 × PIC) on ice for 20 min, washed once with MB 1 buffer, then resuspended in 100 ul MB1 buffer. Mnase concentration for muscle stem cells was predetermined using MNase titration experiments. Chromatin was digested by adding 0.1 μl Mnase (NEB, M0247S) and incubating for 20 min at 37 °C, 1000 rpm. Digestion was stopped by adding 8 μl of 500 mM EGTA and incubating at 65 °C for 10 min. Cells were then washed twice with ice-cold MB2 buffer (10 mM Tris–HCl, pH 7.5, 50 mM NaCl, 10 mM MgCl2). The Mnase-digested DNA ends were polished with T4 PNK (NEB M0201), followed by DNA polymerase Klenow fragment (NEB M0210), and then repaired and labeled by adding biotinylated dATP and dCTP (Jena Bioscience, NU-835-BIO14-S and NU-809-BIOX-S, respectively), and TTP/GTP. After washing with buffer MB3 (50 mM Tris–HCl, 10 mM, MgCl2). Ligation was performed for 4 h at room temperature using T4 DNA ligase (NEB M0202). Dangling ends were removed by a 15-min incubation with Exonuclease III (NEB 0206) at 37 °C. DNAs were de-crosslinked overnight at 65 °C. After DNA extraction by ethanol precipitation, size selection was performed using DNA purification beads to enrich Ligated fragments at about 230 bp. Then, biotin selection was done using 10 μl Dynabeads MyOne Streptavidin C1 beads (Invitrogen 65,001). Libraries were prepared with the NEBNext Ultra II Library Preparation Kit (NEB E7103). Samples were paired-end sequenced (read length = 150 bp) on Illumina’s NovaSeq sequencer.

### Loop interaction frequency analysis

The loops of K562 and HepG2 cells were collected from the Encode dataset (see details in Supplementary Table S9). To remove the confounding effect of genomic distance on interaction frequency, we performed a distance-normalization procedure. First, raw interaction frequencies were log2-transformed after adding a pseudocount of 1 (log2_freq). The linear distance between two interacting loop anchors was calculated as the absolute difference between their genomic coordinates and also log2-transformed (log2_dist). We then fitted a linear regression model with log2_freq as the response variable and log2_dist as the predictor. The residuals obtained from this model were used as distance-corrected interaction frequencies, effectively removing the global distance-dependent trend. To compare the corrected interaction frequencies among different loop types, we performed pairwise Wilcoxon rank-sum tests (Mann-Whitney U tests) for all relevant comparisons. To control the false discovery rate due to multiple testing, the resulting p-values were adjusted using the Benjamini-Hochberg (FDR) method. The corrected frequencies were visualized as boxplots, with pairwise p-values annotated directly on the plot.

### CUT&Tag data analysis

The raw data was first pre-processed by initial quality assessment, adapters trimming, and low-quality filtering and then mapped to the human reference genome (hg38) using *Bowtie2* (v2.3.3.1) [31], and only the non-redundant reads were kept (*Picard* v2.26 was used to remove redundant reads). The peaks (sites) were identified using *MACS2* (v2.2.4) [32]. During the peak calling, the q-value cutoff was set to 0.05 for dG4 and H3K27ac experiments. For the genome location profiling analysis, the genome location of identified peaks was annotated by *ChIPseeker* (v1.38.0) [33]. The reference was set to Homo sapiens based on the UCSC hg38 genome assembly and the known Gene track. And the promoter region was set to upstream and downstream 2000 bps of the transcription start site.

### Micro-C data analysis

The Micro-C raw data was mapped to the mouse reference genome (hg38) using *BWA-MEM* (v0.7.17-r1188) [34]. Next, we used the parse module of the *pairtools* (v1.0.2) [35] pipeline to find ligation junctions in Micro-C libraries. Then, the parsed pairs were then sorted using pairtools sort, and PCR duplication pairs were removed by *pairtools dedup*. Finally, pairtools split was used to generate.pairs files and.bam files. Loops were identified by *HiCCUPS* (v1.22.01) [36] using parameters (-k KR -t 20) and scaled to 5 kb resolution. Cool files were generated by cooler to further compare the interaction difference. Interaction comparisons were calculated by *HiCcompare* (v1.24.0) [37]; “A” value cutoff was set to 15 to filter out interactions with low average expression. Adjusted P value cutoff was set to 0.1 to identify the significant interaction change. As for analysis of TDP-43 and dG4 signal enrichment at loop anchors, the TDP-43 or dG4 ChIP-seq signals of loop anchors was compared with upstream and downstream regions which length equals the anchor length. The test was set to Mann-Whitney U test. As for analysis of the dG4 signal in different type loop anchors, the dG4 ChIP-seq signals of TDP-43+/dG4+ and TDP-43-/dG4+ loop anchors were calculated by mean value. And then these signals were compared by the Mann-Whitney U test.

### dG4 ChIP-seq data analysis

The raw data was first pre-processed by initial quality assessment, adapter trimming, and low-quality filtering by *cutadapt* (v1.15) [38]. Then these trimmed reads were mapped to the human reference genome (hg38) using *bwa mem* (v0.7.17-r1188) [34], and only the non-redundant reads were kept (*Picard* v2.26 was used to remove redundant reads). The peaks (sites) were identified using *MACS2* (v2.2.4) [32]. During the peak calling, the Q-value cutoff was set to 0.05. Only peaks overlapping more than half replicates were considered as high-confidence peaks.

### RNA-seq data analysis

The adapters of raw data were trimmed by *fastp* (v0.23.4) [39]. And the trimmed reads were mapped to the human genome (hg38) by *Hista2* (v2.2.1) [40]. The differential expressed genes were identified by *DESeq2* [41] with parameter setting with adjust p value equals to 0.05 and absolute log2 fold change above 0.5. The FPKM values were quantified by *Cufflinks* (v2.2.1) [42].

### *In vitro* identified PQS data collection

The *in vitro* identified PQS of human were collected from public data [43]. And results (hg19 version) were converted into the hg38 version by *liftOver*s [44].

## Acknowledgements

This work was supported by National Natural Science Foundation of China [82172436 to H.W., 32300703 to X.C.]; Natural Science Foundation of Guangdong Province, China [2024A1515030291 to X.C.]; National Key R&D Program of China [2022YFA0806003 to H.W.]; General Research Fund (GRF) from the Research Grants Council (RGC) of the Hong Kong Special Administrative Region, China [14103522, 14105123, and 14120420 to H.S.; 14100620, 14105823, 14106521, and 14115319 to H.W.]; Theme-based Research Scheme (TRS) from RGC [T13-602/21-N to H.W.]; Strategic Topics Grant (STG) from RGC [STG1/E-403/24-N to H.W.]; Area of Excellence Scheme (AoE) from RGC [AoE/M-402/20 to H.W.]; Health and Medical Research Fund (HMRF) from Health Bureau of the Hong Kong Special Administrative Region, China [10210906 and 08190626 to H.W.]; the research funds from Health@InnoHK program launched by Innovation Technology Commission, the Government of the Hong Kong SAR, China [to H.W.]; Chinese University of Hong Kong (CUHK) Strategic Seed Funding for Collaborative Research Scheme (SSFCRS) [to H.W.].

## Data availability

The dG4 ChIP-seq of K562 and HepG2 cell line data were collected from public dataset (GSE107690 for K562 and HepG2 cell lines). And the TDP-43 ChIP-seq, RNA-seq, ATAC-seq, H3K4me3 ChIP-seq, H3K27ac ChIP-seq, and Hi-C data of K562 and HepG2 cell line data were collected from the ENCODE [45] dataset (Supplementary Table S9). The ChIP-seq data of TFs were collected from the Cistrome Database (Supplementary Table S10) [46]. The raw sequence data and processed data of CUT&Tag and Micro-C of HepG2 cells have been deposited in the Gene Expression Omnibus (GEO) database under accession numbers: CUT&Tag GSE328806; Micro-C, GSE328803.

## Author contributions

F.Y., X.C. and H.W. conceived the research and designed all the experiments; F.Y. performed all the computational analyses; S.Z. performed the CUT&Tag; X.G. performed siRNA transfection and western blot; Y.Q. conducted Micro-C; Y.Z. and H.S. advised bioinformatics analysis; F.Y., X.C. and H.W. wrote the manuscript with input from all authors.

## Inventory of Supplemental Information

### 1. Supplemental figures

Figure S1. Transcriptomic analysis of K562 and HepG2 cells with shNC or shTDP-43 knockdown.

Figure S2. Loop classifications according to TDP-43 binding and dG4 formation.

Figure S3. Analysis of dG4 CUT&Tag in HepG2 cells transfected with siNC or siTDP-43.

### 2. Supplemental tables

Table S1. dG4 peaks identified from public K562 and HepG2 datasets.

Table S2. Genes with promoter TDP-43 binding/dG4 formation in K562 and HepG2 cells.

Table S3. Integrative analysis of public RNA-seq data from K562 and HepG2 cells with shNC or shTDP-43 knockdown.

Table S4. Chromatin loop anchors with or without TDP-43 binding in K562 and HepG2 cells.

Table S5. Chromatin loops identified with public Hi-C datasets from K562 and HepG2 cells.

Table S6. ATAC signal and H3K4me3 signal comparison among different types of loops in K562 and HepG2 cells

Table S7. CUT&Tag data preprocessing and peak identification in HepG2 cells transfected with siNC or siTDP-43.

Table S8. Micro-C data preprocessing and integrative analysis in HepG2 cells transfected with siNC or siTDP-43.

Table S9. Metadata of public RNA-seq, ChIP-seq, and Hi-C data used in this study. Table S10. Transcription factor metadata sourced from the Cistrome database.

Table S11. Sequences of siRNA oligos

### 3. Supplementary figure legends

**Figure S1.**
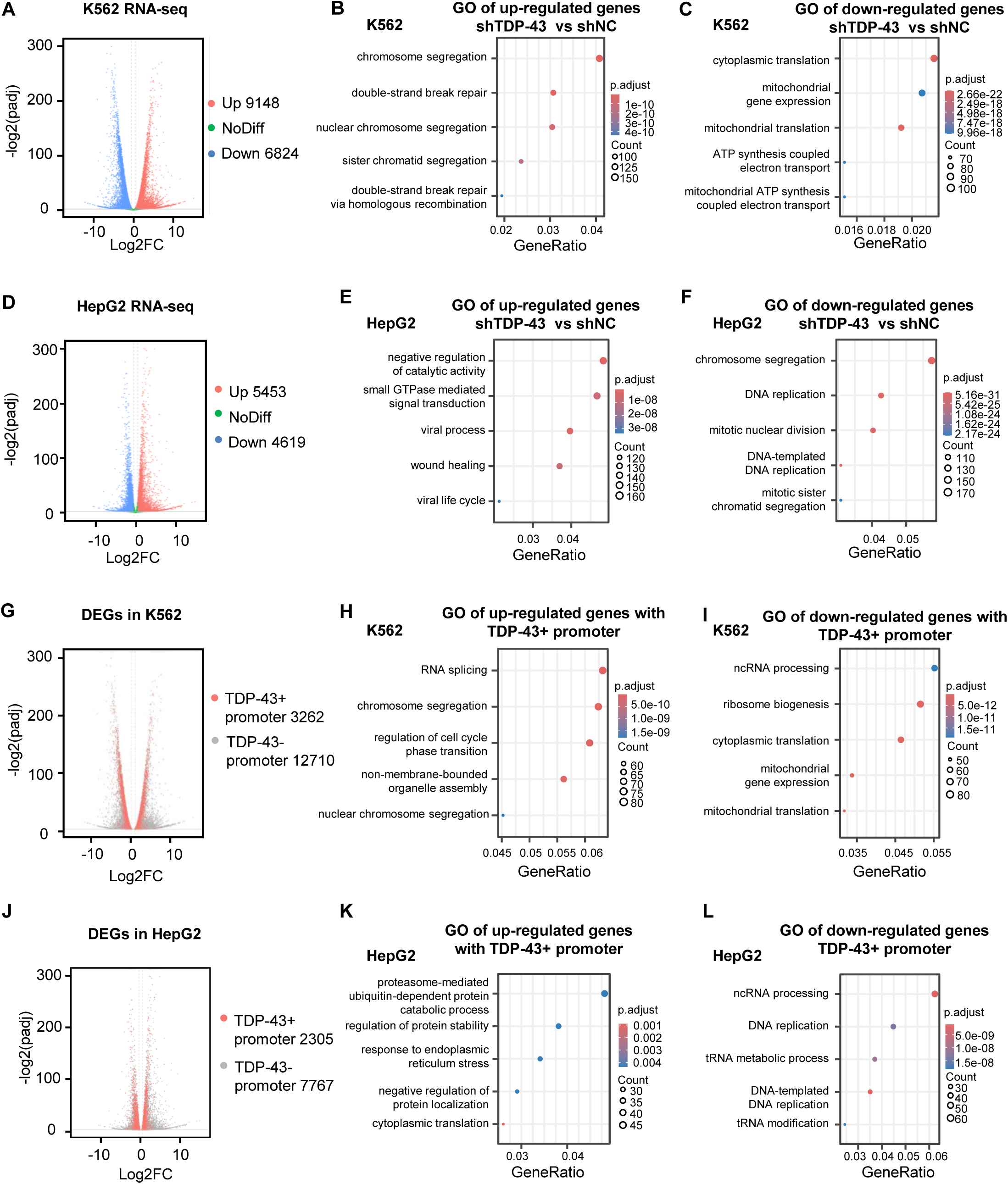
Transcriptomic analysis of K562 and HepG2 cells with shNC or shTDP-43 knockdown. **(A)** Volcano plot showing differentially expressed genes in K562 cells with shTDP-43 vs shNC. **(B)** GO analysis of the up and **(C)** down genes in **(A)**. **(D)** Volcano plot showing differentially expressed genes in HepG2 cells with shTDP-43 vs shNC. **(E)** GO analysis of the up and **(F)** down genes in **(D)**. **(G)** Volcano plot showing differentially expressed genes with or without TDP-43 binding in K562 cells. **(H)** GO analysis of the up and **(I)** down genes with TDP-43 binding in K562 cells with shTDP-43 treatment. **(J)** Volcano plot showing differentially expressed genes with or without TDP-43 binding in HepG2 cells. **(K)** GO analysis of the up and **(L)** down genes with TDP-43 binding in HepG2 cells with shTDP-43.

**Figure S2.**
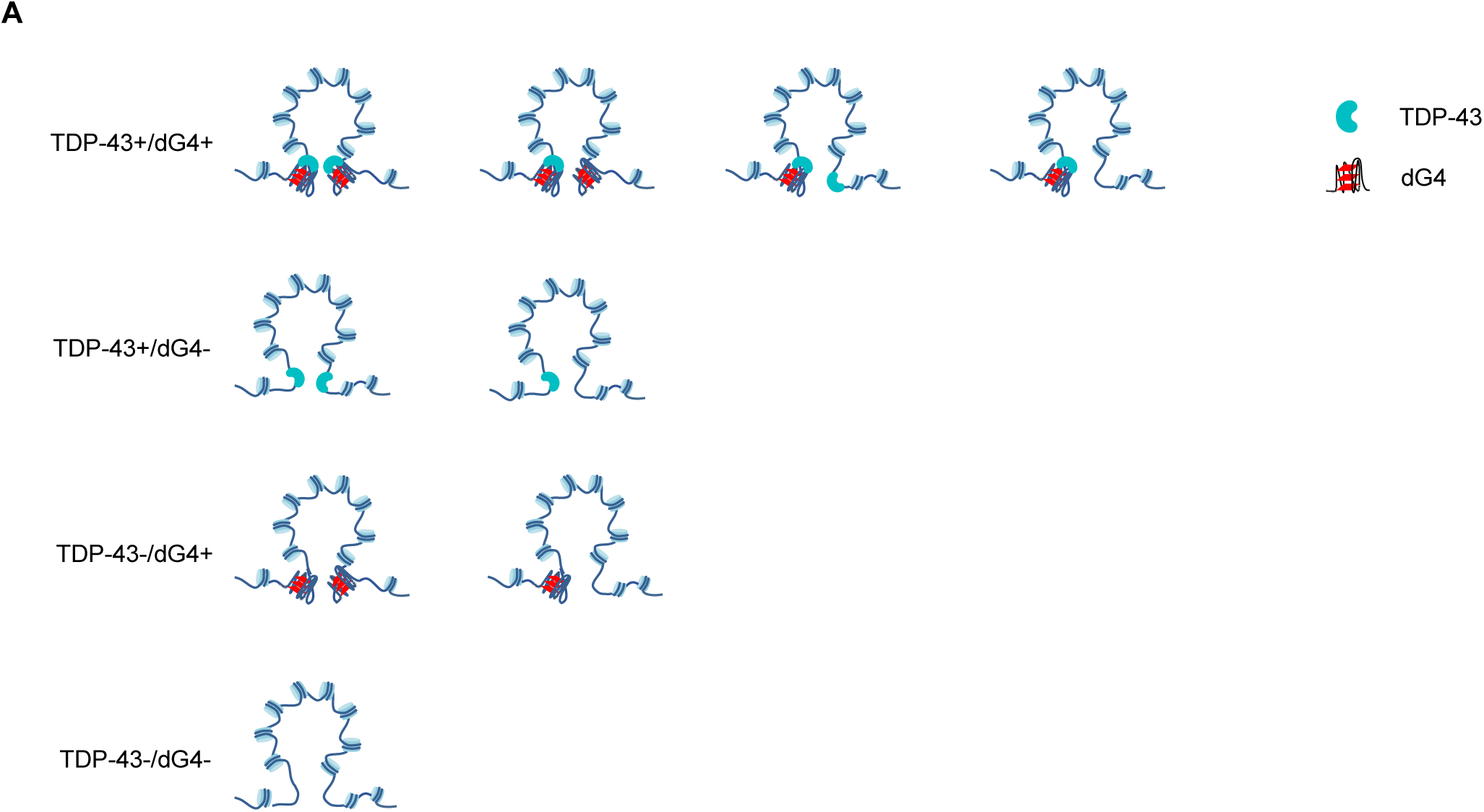
Loop classifications according to TDP-43 binding and dG4 formation. **(A)** Schematic of the 4 types of loops in Fig. 4A.

**Figure S3.**
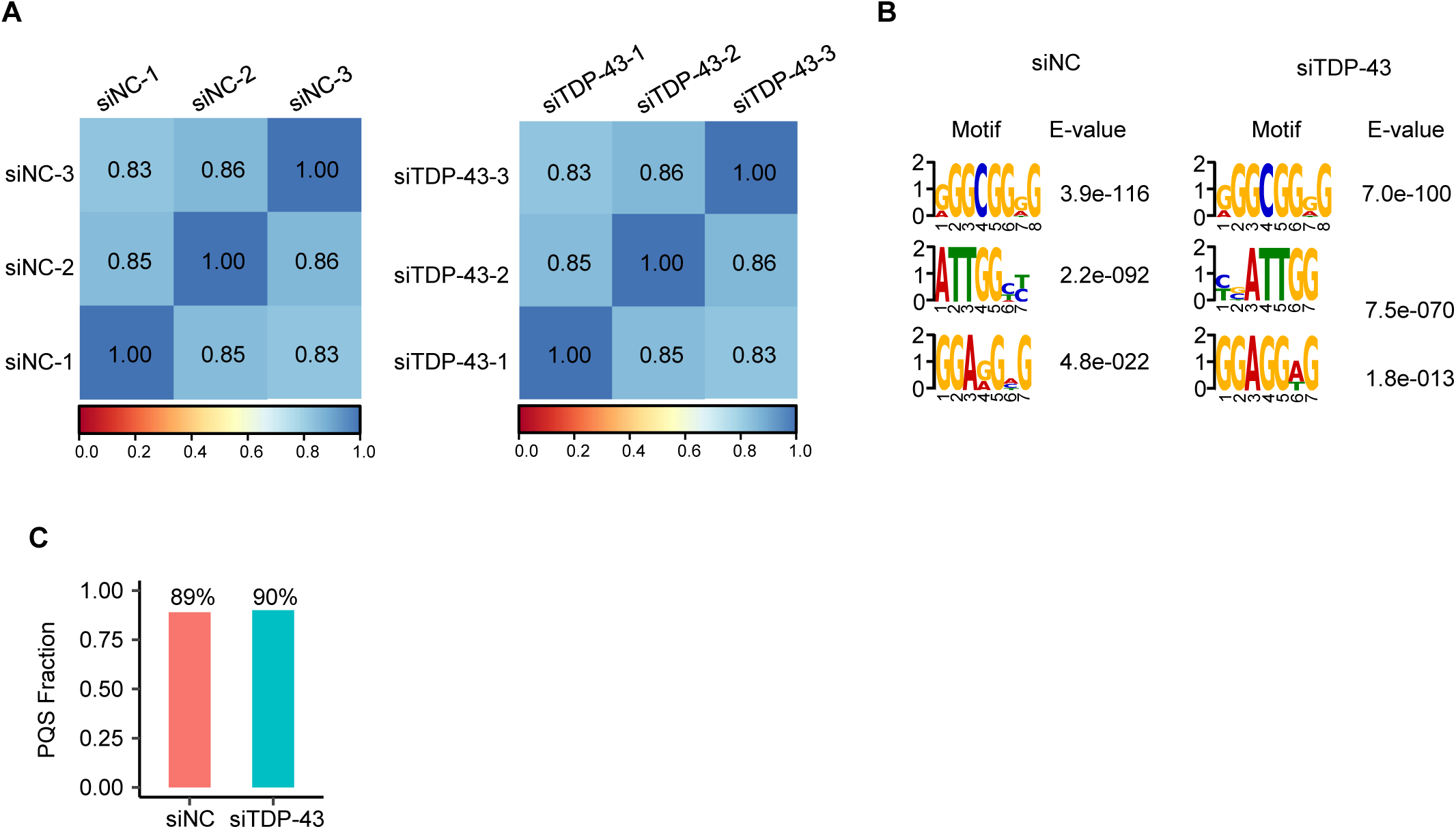
Analysis of dG4 CUT&Tag in HepG2 cells transfected with siNC or siTDP-43. **(A)** Heatmap showing the correlation of dG4 signals in HepG2 cells transfected with siNC or siTDP-43. The Pearson correlation coefficient was calculated. **(B)** Top 3 motifs at dG4 peaks in HepG2 cells transfected with siNC or siTDP-43. **(C)** Proportion of dG4 peaks in HepG2 cells transfected with siNC or siTDP-43 overlapping with PQS.

